# The Ocular Glymphatic Clearance System

**DOI:** 10.1101/2019.12.29.889873

**Authors:** Xiaowei Wang, Nanhong Lou, Allison Eberhardt, Peter Kusk, Qiwu Xu, Benjamin Förstera, Sisi Peng, Yujia Yang, Meng Shi, Anna L. R. Xavier, Ali Ertürk, Richard T. Libby, Lu Chen, Alexander S. Thrane, Maiken Nedergaard

**Author notes:** These authors contributed equally to the work (Xiaowei Wang, Nanhong Lou).

## Abstract

Despite high metabolic activity, the retina and optic nerve head lack traditional lymphatic drainage. We here identified a novel ocular glymphatic clearance route for fluid and wastes via the proximal optic nerve. Amyloid-β (Aβ) was cleared from the vitreous via a pathway driven by the ocular-cranial pressure difference. After traversing the lamina barrier, intra-axonal Aβ was cleared via the perivenous space and subsequently drained to lymphatic vessels. Light-induced pupil constriction enhanced, while atropine or raising intracranial pressure blocked efflux. In two distinct murine models of glaucoma, Aβ leaked from the eye via defects in the lamina barrier instead of directional axonal efflux. The discovery of a novel pathway for removal of fluid and metabolites from the intraocular space prompts a reevaluation of the core principles governing eye physiology and provides a framework for new therapeutic approaches to treat common eye diseases, including glaucoma.

**One Sentence Summary:** Glymphatic pathway clears ocular amyloid-β via optic nerve and fails in glaucoma.

## Main Text

Like the brain inside the cranial vault, the internal structures of the eye are contained within a confined space, which necessitates tight control of fluid homeostasis. Yet, both the eye and brain are largely devoid of traditional lymphatic vessels, which are critical for the clearance of fluid and solutes from peripheral tissues (*1*, *2*). Recent discoveries have shown that the brain possesses a quasi-lymphatic system, termed the glymphatic system (*3*), and that traditional lymphatic vessels are also present in the dura mater, one of three layers of fibrous membranes lining the exterior surface of the brain (*4*, *5*). In the brain, glymphatic/lymphatic transport contributes to clearing of amyloid-β (Aβ), a derivative of amyloid precursor protein (APP) that is one of the main constituents of amyloid plaques in the brain (*6*) and likewise of amyloid deposits in the retina (*7*). In the eye, several hypothesis-driven investigations have pointed to the existence of an ocular glymphatic system (*8*–*10*). Experimental data have supported this concept by demonstrating retrograde cerebrospinal fluid (CSF) transport along the perivascular spaces in the optic nerve (*11*). Since the highly metabolically active neural tissues of the retina produce Aβ (*3*, *12*) and other potentially neurotoxic protein cleavage products like tau (*13*), we explicitly sought evidence of an intraocular anterograde glymphatic/lymphatic clearance system. First, we injected HiLyte-594-tagged human amyloid-β (hAβ) into the vitreous body of mice and visualized tracer distribution one hour later. Then we used uDISCO whole-body clearing (*14*) to analyze the eye and tracer in see-through mouse heads. Surprisingly, three-dimensional reconstruction of light-sheet microscopy data revealed that hAβ tracer exited the eye along the optic nerve (**Fig. 1A, fig. S1A**). Transport along the optic nerve also occurred following non-invasive intravascular delivery of either the radiolabeled K^+^-analogue (^86^Rb) or FITC-cadaverine, a tracer that is permeable to the retinal-blood barrier, but not the blood-brain barrier (*15*, *16*) (**fig. S1B-D**). Whole-mount preparation of the optic nerve confirmed that hAβ tracer is transported anterograde along the optic nerve (**Fig. 1B**). Moreover, use of reporter mice in which arteries and arterioles can be identified by DsRed expression in mural cells revealed that hAβ tracer preferentially accumulated in the perivascular space along the optic nerve veins, rather than along arterioles (**Fig. 1C**). In contrast to hAβ tracer, Alexa Fluor (AF)-dextran (3, 10, and 500 kDa) did not enter the optic nerve after intravitreal administration (**Fig. 1D, fig. S1E**). Sectioning and high-resolution imaging of the optic nerve showed that in addition to perivenous accumulation, hAβ tracer was also transported along neuron-specific Class III β-tubulin (TUJ1)-positive axons, consistent with rapid neuronal uptake of hAβ (*17*) (**Fig. 1 E-F**). Glial cells of the oligodendrocytic (*Olig2*^+^) and the astrocytic (*Glt-1*^+^) lineage also exhibited sparse hAβ tracer uptake (**fig. S2A-C**). The hAβ tracer accumulation peaked 260 ± 17 μm (mean ± SEM) from the optic nerve head and tapered off gradually in the distal nerve, suggesting that hAβ tracer must exit the optic nerve anterior to the optic chiasm. This notion was supported by the observation that the dural sheet surrounding the proximal segment of the optic nerve densely accumulated tracer (**Fig. 1E**). Additional analysis revealed traditional lymphatic vessels that were embedded in the dural sheath and surrounding loose tissue (**fig. S2D-F**). These vessels did not contain red blood cells or stain with intravascularly-delivered lectin, but they labeled positive for the conventional lymphatic markers, lymphatic vessel endothelial hyaluronan receptor 1 (LYVE-1), vascular endothelial growth factor receptor 3 (VEGFR3) and podoplanin (PDPN) (*2*, *5*). Imaging after dual injections of hAβ tracer in the vitreous body in conjunction with AF-dextran (10 kDa) in the cisterna magna (CM) highlighted that the tracers were transported within the optic nerve in both anterograde and retrograde directions with limited spatial overlap (**Fig. 1H-I**). As reported previously, tracers injected in the CM were transporter along perivascular spaces (*11*). Use of reporter mice with DsRed expression in mural cells revealed that CM-injected tracers were predominantly transported along the periarterial and pericapillary spaces (**Fig. 1J, fig. S2H**), in contrast to the intravitreal tracers, which accumulated along the veins (**Fig. 1C**). Collectively, these observations show that hAβ is transported by axons and along the perivenous space in the metabolically vulnerable (unmyelinated) initial segment of the optic nerve (*18*) after intravitreal delivery. From there, hAβ exits the optic nerve via dural lymph vessels located in the outer layer of meninges lining the optic nerve and orbital lymphatics, as recently reported (*19*). In support of this model, hAβ had accumulated in the ipsilateral cervical lymph nodes in animals examined three hours after intravitreal injection (*2*) (**Fig. 1G, fig. S2G**).

**Figure 1.**
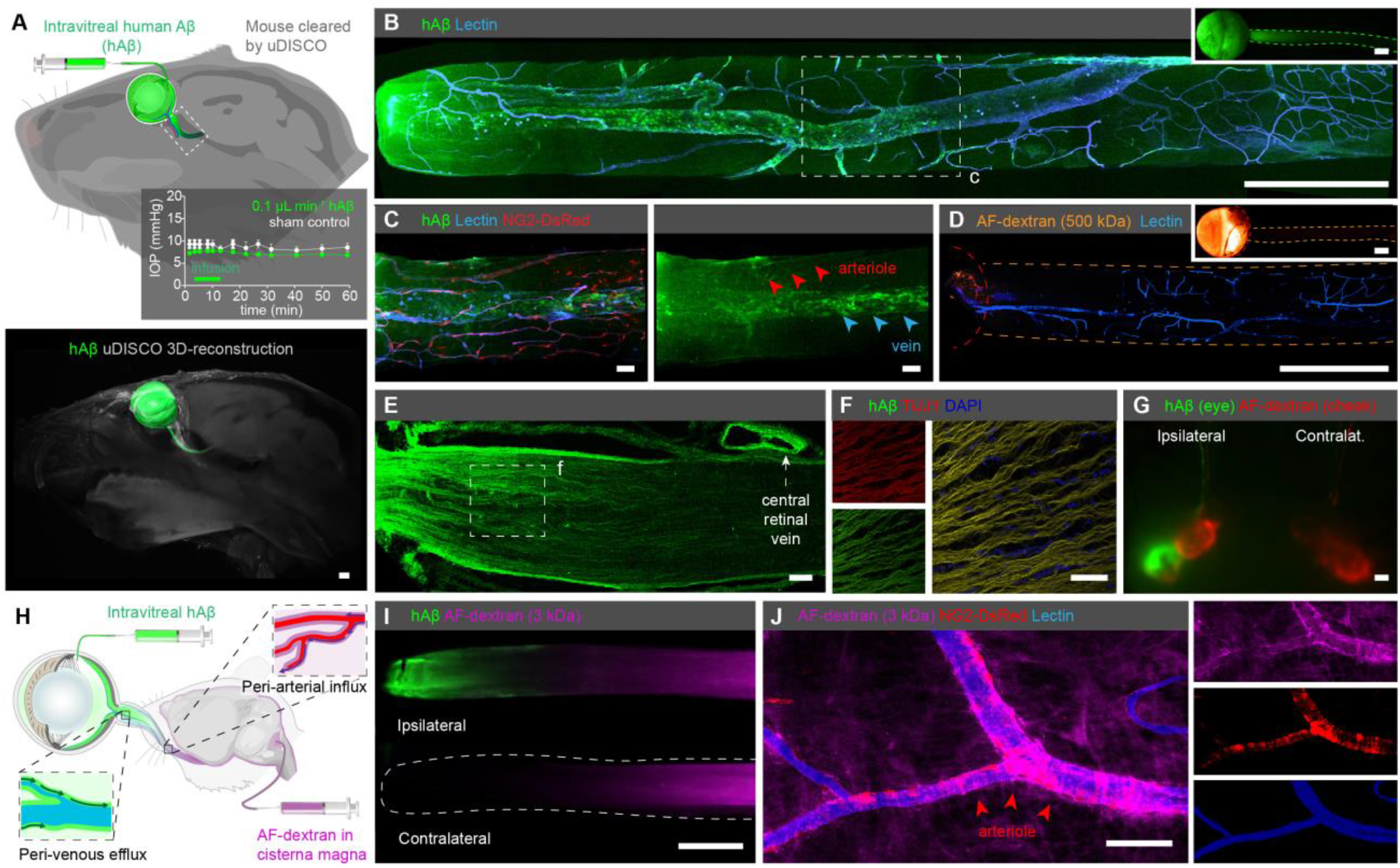
Existence of an ocular glymphatic clearance system. **(A)** Upper-panel: schematic of intravitreal injection of hAβ. Insert: IOP during injection (mean±SEM, n=5, *p*=0.0450-0.5970, unpaired two-tailed *t*-test). Rectangle indicates proximal optic nerve displayed in panels **B-J**. Lower-panel: uDISCO-cleared transparent mouse heads one hour after hAβ injection shows anterograde-transport of hAβ along the optic nerve. (**B**) Confocal of ipsilateral optic nerve following injection shows hAβ accumulation along the central retinal vein. **B** and **D** inserts display macroscopic images of the eye and optic nerve injected with respective tracers without background subtraction. (**C**) Reporter mouse with DsRed-tagged mural cells (vascular smooth muscle cells and pericytes) shows that hAβ accumulates around optic nerve veins rather than arterioles 30 min after injection. (**D**) AF-dextran does not pass the glial-lamina but is retained within the eye 30 min after intravitreal injection. (**E-F**) Confocal show tracer transported along TUJ1-positive axons. Note tracer accumulation in the dural lining of the nerve. (**G**) Cervical lymph nodes exhibiting intense hAβ labeling three hours after injection. (**H**) Schematic of the double injections. (**I**) Double injections in the vitreous body (hAβ) and cisterna magna (AF-dextran) show that the tracers are transported in both antero- and retrograde manners. (**J**) The retrograde-transport of AF-dextran occurred primarily along the periarterial space. (scales **A, B, D, G**: 500μm; **C, E, F, I, J**: 50μm).

Pressure difference is a principal driving force of directional fluid transport. Since IOP under physiological conditions exceeds intracranial pressure (ICP), it is possible that the pressure difference across the lamina cribrosa contributes to fluid flow along the optic nerve (*20*). To define the role of the translaminar pressure difference in transport of hAβ tracer, we manipulated ICP by either withdrawal or injection of artificial CSF in the CM while recording ICP (**Fig. 2A**). Decreasing intracranial pressure from the baseline level of 3.6 ± 0.4 mmHg to 0.5 ± 0.2 mmHg was linked to a sharp increase in the total hAβ tracer signal and the peak intensity of hAβ tracer transport in the proximal optic nerve 30 min after intravitreal injection (**Fig. 2B-E**). Conversely, increasing ICP to 16.7 ± 0.6 mmHg (**Fig. 2A**) blocked hAβ tracer transport in the optic nerves (**Fig. 2B-E**). These data show that hAβ tracer transport along the optic nerve is sensitive to manipulation of the translaminar pressure gradient.

**Figure 2.**
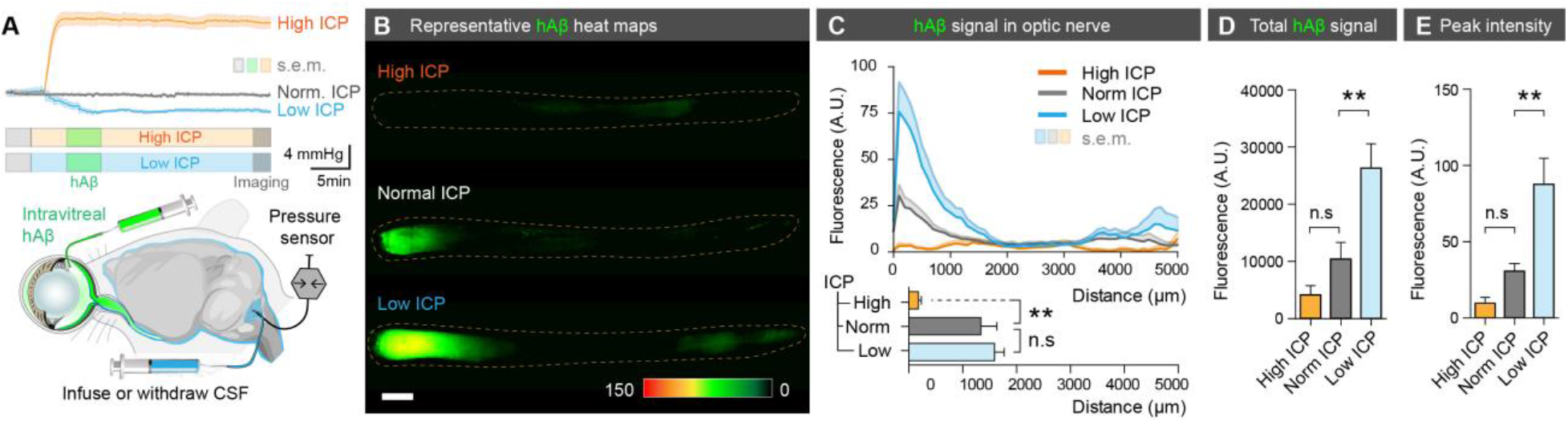
The translaminar pressure difference drives ocular glymphatic outflow. (**A**) Schematic of the setup used for analyzing hAβ transport following intravitreal injection while manipulating ICP. Upper-panel: mean ICP (±SEM) plotted as a function of time in the high, normal, and low ICP groups (n =10-12). (**B**) Representative background-subtracted heat-maps of hAβ in the optic nerve from high, normal, and low ICP groups. (**C**) Upper-panel: averaged fluorescent intensity profiles of hAβ in the optic nerves from the three groups. Lower-panel: the distance of tracer transport (mean±SEM, n=6, \*\**p*<0.01, n.s. *p*=0.5929 for distance, 0.2858 for total signal, 0.2924 for peak intensity, one-way ANOVA followed by Dunn’s *post hoc* test). (**D-E**) Total hAβ signal and peak intensities in the optic nerve 30 min after intravitreal injection (mean±SEM, n=6 for each group, \*\**p*<0.01, one-way ANOVA followed by Dunn’s *post hoc* test). (scale: 500μm).

Prior observations have shown that repeated constriction of the pupil and ciliary body during accommodation enhances aqueous outflow in healthy human subjects, hypothesized to be via trabecular and uveoscleral routes (*21*, *22*). We next asked whether the transport of hAβ along the optic nerve is influenced by physiological pupil constriction *in vivo*. To address this question, we compared transport of hAβ tracer in mice stimulated with light at 1 Hz to control mice kept in darkness. A subset of mice exposed to light stimulation received atropine (1%, eye drops) to block the light-induced pupillary constriction reflex, while another subset of mice received pilocarpine (2%, eye drops) to achieve static constriction without light stimulation (**Fig. 3A**). The analysis showed that light stimulation increased hAβ transport in the optic nerve. The total hAβ tracer signal, peak intensity, and distance of transport were all sharply increased by light stimulation at 30 min after hAβ administration (**Fig. 3B-D**). Tracer accumulation in mice kept in darkness increased slowly and reached the same amplitude as that in the light-stimulated eyes one to two hours after intravitreal injection of hAβ tracer (**fig. S4A-I**; the findings were also reproduced in rats: **fig. S3A-E**). Infrared pupillometry confirmed that the light stimulation induced alternating constriction and dilation of the pupil, detected as high variance of pupil size. In contrast, animals kept in the dark, as well as light-stimulated animals pretreated with atropine, exhibited essentially no pupil movement (**Fig. 3E-G**). Atropine completely blocked the light-induced pupil constriction, as well as the light-related acceleration of hAβ tracer transport (**Fig. 3 B-G**), while pilocarpine-induced tonic pupillary constriction did not appear to affect transport (**Fig 3 B-D**). Intravitreal hAβ tracer was not transported postmortem (**Fig. 3B**, **fig. S4B and F**), suggesting that passive diffusion only insignificantly contributes to hAβ tracer dispersion along the optic nerve. These data suggest that repeated pupil and potentially ciliary body constriction propels intraocular fluid dispersion along the optic nerve, resulting in enhanced hAβ tracer transport. Of note, combining light stimulation with manipulations of ICP did not change net hAβ tracer transport compared with either low or high ICP alone (**fig. S5A-E**). Thus, sufficiently large pressure changes can override the effect of natural light stimulation.

**Figure 3.**
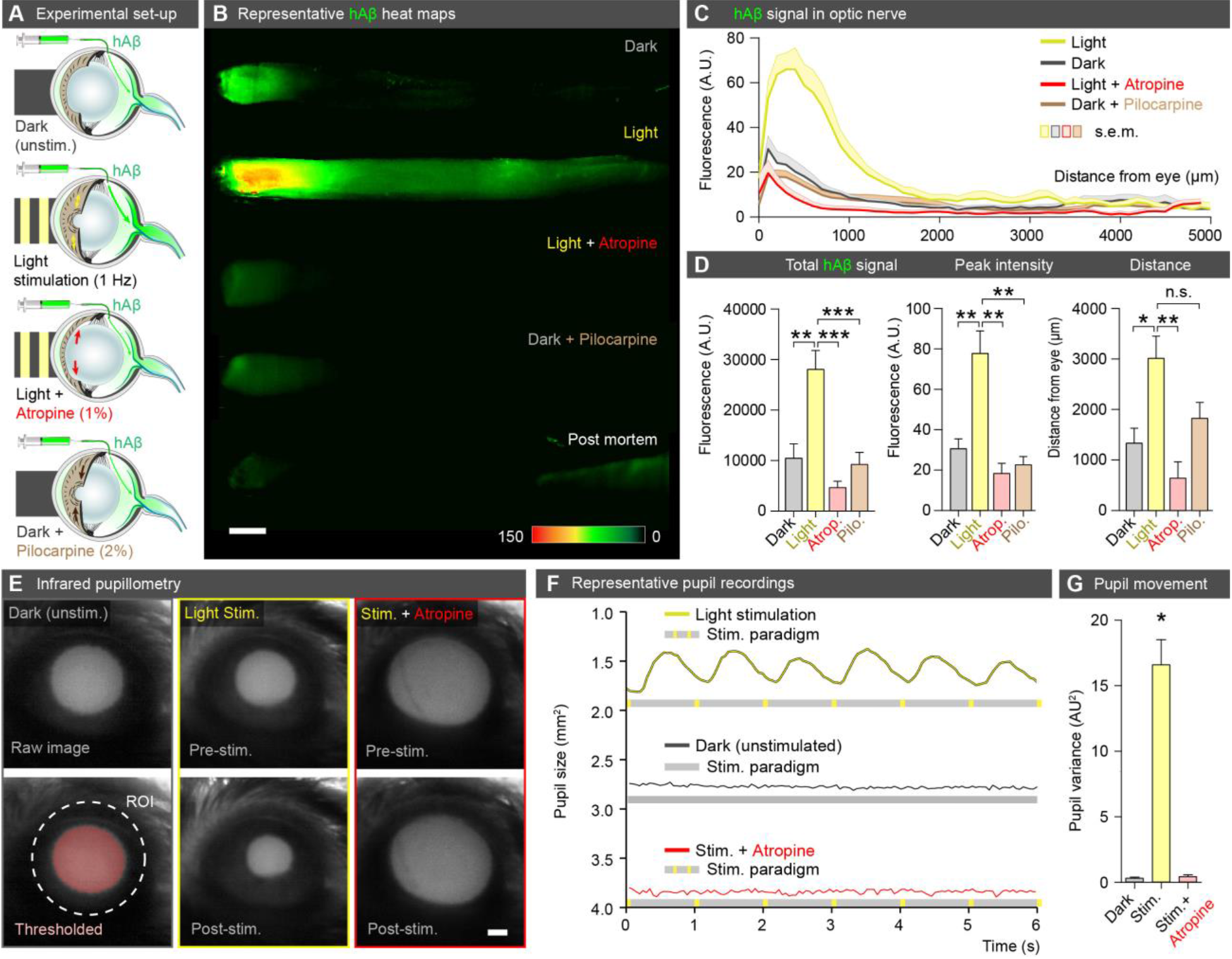
Light stimulation enhances hAβ along the optic nerve. (**A**) Schematic of the experimental groups. The first group was kept in darkness. The second group was exposed to 1 Hz light-stimulation (100 ms duration, 5 lumens). The third group was pretreated with atropine (1%) before exposure to 1 Hz light-stimulation. The fourth group was pretreated with pilocarpine (2%) and kept in darkness. (**B**) Representative background-subtracted heat-maps of optic nerves from the four groups 30 min and a postmortem group 120 min after injection of hAβ. (**C**) Averaged fluorescent intensity profiles of optic nerves from the four groups (mean±SEM, n=6-19). (**D**) hAβ signal mapped as total signal, peak intensity, and distance of the hAβ transport (mean±SEM, n =6-19, \**p*<0.05, \*\**p*<0.01, \*\*\**p*<0.001, n.s. *p*=0.0756, one-way ANOVA followed by Dunnett’s *post hoc* test). (**E**) Infrared pupillometry tracking of the pupil size and light-induced constriction with and without atropine pre-treatment. The pupil area (mm^2^) was determined by auto-thresholding. (**F**) Representative pupillometry recordings in dark-exposed and in light-stimulated mice with and without atropine administration. (**G**) Comparison of the variance of pupil area in these groups (mean±SEM, n=3, unpaired two-tailed *t*-test, \**p*<0.05). (scale: 500μm).

Glaucoma is a group of diseases characterized by progressive and irreversible injury to the optic nerve head and retinal ganglion cell (RGC) degeneration, leading to blindness. Increased IOP is a leading risk factor for glaucoma. Our present observations raised the question of whether glaucoma is linked to pathological changes in ocular glymphatic solute transport. To test this, we undertook tracer studies in two distinct murine models of glaucoma and chronic ocular hypertension based on two separate mouse strains. DBA/2J mice develop a depigmenting iris disease that leads to age-related ocular hypertension (*23*) (**Fig. 4A**). Chronic IOP elevation results in significant RGC loss and glaucomatous optic nerve degeneration in the majority of aged eyes from DBA/2J mice (*24*) (**fig. S7E and F**), but rectifying their IOP with pharmacological, genetic, or surgical interventions significantly ameliorates RGC death (*25*). The chronic circumlimbal suture (CLS) model employs oculopression to reduce aqueous drainage, increase IOP over a 1-month period, and secondarily cause RGC loss in non-pigmented CD-1 mice (*26*). Our analysis of young DBA/2J mice showed that neither IOP nor hAβ tracer transport along the optic nerve differed from that in the D2-Gpnmb^+^ (D2-control) mice, a genetically-matched control strain that does not develop ocular hypertension (**Fig. 4A and B, fig S7D**). About 50% of DBA/2J mice are severely affected by glaucoma at 11 months, but IOP is usually normalized or only mildly elevated at this stage of the disease and thus less likely to directly influence tracer outflow (*24*). Surprisingly, using whole-mounts of optic nerve, we found that hAβ tracer transport was sharply increased in an equivalent subset of 11-month-old DBA/2J mice (**Fig. 4A-D, fig. S6A-D**), with total hAβ tracer signal significantly increased in 11-month-old DBA/2J mice compared with age-matched D2-control mice (**Fig. 4B-D)**. Similarly, we found that CD-1 CLS but not sham control mice exhibited significantly increased hAβ tracer transport one month after surgery, when IOP had normalized (*26*) (**Fig. 4B-D, fig. S6E**). High-resolution confocal imaging of 11-month-old DBA/2J optic nerves revealed that hAβ tracer was located primarily in the perivascular space, or outside the RGC axons, as opposed to the intra-axonal distribution noted in age-matched D2-control and young DBA/2J mice (**Fig. 1F, fig. S7A, fig. S6B**). This prompted us to ask whether the glial lamina normally acts as a high-resistance barrier that hinders transport of macromolecules. If so, then a corollary of this proposition would be that a failure of the lamina barrier might be one of the hallmarks of glaucoma (*27*, *28*). To test this dual hypothesis, we assessed the passage of increasing molecular sizes of AF-dextran across the lamina following intravitreal injection in D2-control and DBA/2J mice (3, 10, 500 kDa), as well as CLS and sham control CD-1 mice (500 kDa). We found that even the smallest dextran (3 kDa, compared to 4.3 kDa for hAβ40) failed to pass the glial lamina in the D2 and CD-1 control mice (**Fig. 1D, Fig. 4E, fig. S6G-L**). In contrast, we saw significant dispersion of all molecular sizes of dextran in 11-month-old DBA/2J and CLS mice, with tracer being detectable several mm distal to the lamina (**Fig. 4E-H, fig. S6B-L**). Confocal imaging confirmed that the dextrans were present in the perivascular space, or outside the few remaining axons in 11-month-old DBA/2J mice (**fig. S7B-D, fig. S6B**). These observations suggest that the lamina barrier in the healthy eye diverts extracellular fluid into the axonal compartment at the optic nerve head and thus facilitates directional axonal fluid transport. Although our data do not aim to address the proximal cause of lamina barrier failure, they show that openings in the barrier divert flow of the ocular fluid in the optic nerve from the axonal to the extracellular compartment. Ultrastructural analysis revealed large defects in the glial barrier in old DBA/2J mice, as previously reported (*27*, *29*, *30*) (**Fig. 4I, fig. S7G**). We also tested the alternative hypothesis that there could be a general expansion of the extracellular space volume fraction in response to axonal loss in old DBA/2J mice. Such an increase in extracellular space of the optic nerve would be expected to facilitate pressure driven-hAβ transport. However, real-time TMA^+^ electrophoretic analysis failed to confirm this hypothesis; the mean volume fraction (α) was ~10% of the total volume and did not differ between D2-control mice and DBA/2J mice (**Fig. 4J-K**). The tortuosity factor (λ) exhibited a non-significant trend toward reduction in old DBA/2J, perhaps reflecting axonal loss (*31*) (**Fig. 4J**).

**Figure 4.**
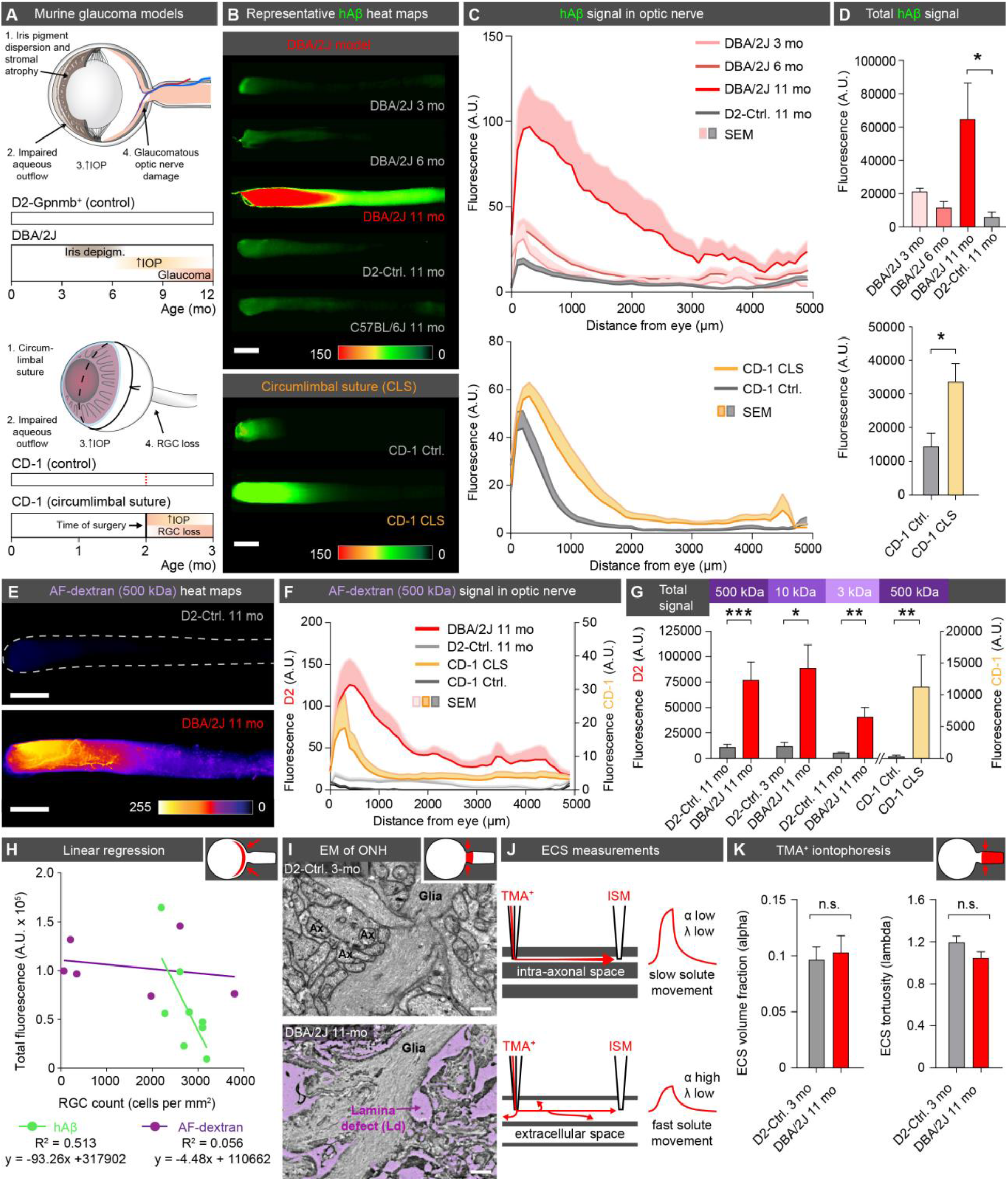
Disruption of the lamina barrier in two distinct murine models of glaucoma reveals a redirection and pathological enhancement of ocular glymphatic outflow. (**A**) Schematic of disease progression in the DBA/2J strain and the chronic CLS model (CD-1) (*24*, *26*). (**B**) Representative background-subtracted heat-maps of optic nerves 30 min after hAβ injection in young, middle-aged, and old DBA/2J mice, and old D2-control mice, as well as CD-1 CLS and CD-1 control mice. (**C**) Averaged fluorescent intensity profile of hAβ distribution along the optic nerve in old DBA/2J or CD-1-CLS compared to respective controls (mean±SEM, n=6-12). (**D**) Total hAβ signal in old DBA/2J or CD-1-CLS compared to respective controls (mean±SEM, n=6-12, \**p*<0.05, Kruskal-Wallis followed by Dunn’s *post hoc* test for DBA/2J model, unpaired two-tailed *t*-test for CLS model). (**E**) Representative background-subtracted heat-maps of optic nerves from old DBA/2J and D2-control mice 30 min after intravitreal administration of AF-dextran. (**F**) Averaged fluorescent intensity profile of AF-dextran along the optic nerve in old DBA/2J or CD-1-CLS compared to respective controls (mean±SEM, n=6-9). (**G**) Total signal of different sized AF-dextrans in optic nerve from old DBA/2J or CD-1 CLS compared to respective controls (mean±SEM, n =4-9, \**p*<0.05, \*\**p*<0.01, \*\*\**p*<0.001, unpaired two-tailed *t*-test or Mann-Whitney test). (**H**) Total hAβ or AF-dextran signal in the optic nerves of old DBA/2J mice plotted as a function of RGC density in their retinas (n=6-8). (**I**) Electron micrographs of the glial lamina region from young D2-control and old DBA/2J mice. (**J**) Schematic of real-time TMA^+^ iontophoresis measurement. (**K**) TMA^+^ measurements of α (extracellular volume space) and λ (extracellular tortuosity) (mean±SEM, n=6-20, *p*=0.765 for α, 0.177 for λ, unpaired two-tailed *t*-test) (scale **B**, **E**: 500μm; **I**: 0.5μm).

The demonstration of an ocular glymphatic clearance system driven by the translaminar pressure gradient raises several questions. For example, is the ocular glymphatic system the principal clearance route for protein metabolites such as Aβ produced by RGCs and other retinal cells? Are any subtypes of RGCs or other retinal cells more important for glymphatic transport? Does solute elimination by this pathway occur preferentially during daytime, driven by pupil constriction, or even by general (saccadic) eye movement, and would this explain why ageing and diseases affecting the iris/pupil/ciliary body are so closely linked to glaucoma (*32*, *33*)? Do physical defects in the glial lamina redirect physiological intra-axonal transport to lower resistance extracellular and perivascular pathways, resulting in reduced axonal transport of endogenous APP and its cleavage product, Aβ (**fig. S8A, Movie S1**)? Interestingly, a build-up of amyloid and other wastes in a so-called optic nerve head drüsen is found in 2-4% of the general population and more frequently in patients with narrow optic canals (*34*). An immunohistochemical analysis of APP showed that the protein indeed accumulated in axons of old DBA/2J mice, in accordance with the literature (*35*) (**fig. S8B**). It is therefore possible that abnormal transport of APP/Aβ or other metabolites driven by the increase in translaminar pressure contributes to the degeneration of RGC axons at the early stages of glaucoma or following optic nerve injury (*36*, *37*). Also, we predict that the pathological increase in extracellular ocular fluid fluxes along the optic nerve in later-stage glaucoma triggers additional stress in the form of both heightened metabolic demands and loss of essential cytosolic components, such as vitamin B_3_ (*38*). Conversely, our data suggest that the increased ICP and papilloedema of the optic nerve head observed in patients with idiopathic intracranial hypertension and in astronauts after long-duration in microgravity (*39*, *40*) would directly inhibit ocular glymphatic transport. Taken together, our analysis provides the first evidence for the existence of a highly polarized ocular glymphatic clearance system, an entirely new drainage pathway of the eye with profound implications for our understanding of eye health and disease.

## Supporting information

supplemental materials for The Ocular Glymphatic Clearance System

## Acknowledgements

We thank Karen L. Bentley and Gayle Schneider from URMC Electron Microscope Research Core Facility, and Wei Song and Jinwook Jung for technical support; Charles Nicholson for discussions, Vinita Rangroo Thrane and Paul Cumming for comments on the manuscript, and Dan Xue for graphic illustrations and animations.

## Author contributions

X.W., N.L., A.S.T. and M.N. designed the experiments, performed the data analysis, prepared the figures and wrote the manuscript. B.F. and X.W. performed uDISCO experiments. Q.X. performed TMA^+^ recordings. X.W. and P.K.J. performed pupillometry. N.L. performed IOP recordings. X.W., N.L., A.E., Q.X., S.P., A.L.R.X. performed the remainder of imaging and data collection. X.W., L.C., Y.Y., M.S., N.L., and R.T.L. generated murine models. Data needed to evaluate conclusion is presented in this manuscript or supplementary material.

## Supplementary Materials

Materials and Methods

Figs. S1 to S8

References and notes

Captions for Movies S1

Movie S1

