## supplemental materials for The Ocular Glymphatic Clearance System for "The Ocular Glymphatic Clearance System"

**This PDF file includes:**

Materials and Methods
Figs. S1 to S8
References and notes

### Materials and Methods

#### Experimental animals

Adult 2-12-month-old C57BL/6J, NG2-DsRed, *Glt-1*-eGFP, FVB, DBA/2J, DBA/2J-*Gpnmb*$^{+}$/SjJ and CD-1 mice of either sex (Jackson Laboratory), as well as two-month-old Brown Norway rats (Charles River Laboratories) were used. All rodents were bred and housed in standard housing conditions at the Universities of Copenhagen, Rochester, or California, Berkeley in temperature-controlled rooms at 24°C with a normal 12-hour light/dark cycle. The animals had free access to food and water. Age-matched mice from both C57BL/6J and DBA/2J-*Gpnmb*$^{+}$/SjJ (D2-controls) strains were used as controls whenever possible for the DBA/2J strain. Our analysis showed no difference between the two strains at a young or old age with regard to hAβ tracer transport, including total hAβ tracer signal and peak intensity from both the dark, unstimulated group and light-stimulated group; TMA analysis, and IOP ($p$=0.16-0.87, Mann-Whitney test or unpaired two-tailed $t$-test). We did not observe any difference between C57 and D2-control mice with regard to the small dextran (3 kDa) transport, including total tracer signal and peak intensity ($p$=0.12-0.80), indicating the integrity of the glial lamina is comparable between these two groups of mice. The circumlimbal suture (CLS) model was adapted from Liu et al. (*26*). Briefly, mice were anesthetized as described below, and a suture was passed transconjunctivally around the equator of the eye, with suture tension adjusted to achieve the desired IOP increase. Mice with initial IOP > 55 mmHg were excluded from the analysis, as this level of IOP can impair retinal perfusion (*26*). Age-matched CD-1 mice without the surgical procedure were used as controls for the CLS model. All procedures were approved and in accordance with the guidelines of institutional Animal Care and Use Committee and the National Institutes of Health Guide for the Care and Use of Laboratory Animals at all universities. The experiments were designed to minimize the number of animals utilized.

#### *In vivo* intravitreal and intracisternal tracer injection

All rodents were anesthetized with ketamine/xylazine cocktail (100 mg/kg and 20 mg/kg, respectively). Once reflexes had ceased, mice or rats were mounted in a stereotaxic head frame, and body temperature was maintained by a heating pad (Harvard Apparatus). A conjunctival peritomy was performed to expose the sclera at 11 to 1 o’clock. A 34G needle connected to PE20 polyethylene glycol tubing and a Hamilton syringe was inserted into the vitreous chamber 1-2 mm posterior to the corneoscleral junction at a depth of approximately 1-2 mm. To map lymphatic drainage three hours following intravitreal tracer injection of hAβ-647, mice were imaged with a stereofluorescent microscope (MVX10, Olympus) in the supine position. The cervical region was exposed before imaging. We pre-labeled mouse cervical lymph nodes by bilateral injection of fluorescein into the mouse cheek as described (*19*). To compare the performance of the ocular glymphatic system in different physiological conditions, 1 μl of one of the following tracers was injected over a period of either five or ten min by a syringe pump (11 Plus, Harvard Apparatus): HiLyte Fluor 488-, HiLyte Fluor 555-, or HiLyte Fluor 647-conjugated hAβ (1-40, AnaSpec) FITC-conjugated-cadaverine (Thermo Fisher Scientific), FITC-, Texas Red-, Alexa Fluor (AF) 488- or AF 555-

conjugated dextrans (3 kDa, 10 kDa, 500 kDa, referred to as AF-dextran for simplicity) (Thermo Fisher Scientific). (**Fig.1a**). We saw no statistical differences in hAβ tracer distribution between the five- and ten-minute injections with respect to fluorescent signal and peak intensity (n=6-8, $p$=0.527-0.649, unpaired two-tailed $t$-test). We used a needle holder attached to a hand manipulator (WPI) to secure the needle in position for the duration of the experiment. An eye lubricant (GenTeal Lubricant Eye Gel, Alcon) was applied to the cornea during the experiment. Photopic flicker (1 Hz, 100 ms, 5 lumens) from an LED light source (Pelican L4) was used to stimulate the tracer-injected eye under control of a Master 8 stimulator (A.M.P.I.). A subset of 1 Hz light-stimulated mice was treated with 1% atropine (Sigma-Aldrich) by topical instillation to the stimulated eye. A subset of dark, unstimulated mice was treated with 2% pilocarpine (Sigma-Aldrich) by topical instillation. During the experiment, the pupil sizes of subsets of mice from the dark, unstimulated group, 1 Hz light-stimulated group, and atropine-treated group were tracked by an infrared camera (DCC3240N, Thorlabs). Intracisternal (cisterna magna, CM) injections were performed in anesthetized mice fixed in a stereotaxic frame. A 30G needle was inserted into the CM, and 0.5% AF-dextran (Thermo Fisher Scientific) was infused at a rate of 1.5 µl/min over 10 min with a syringe pump (Harvard Apparatus).

**Ultimate 3D imaging of Solvent-Cleared Organs (uDISCO)**
One hour following intravitreal injection of hAβ-HiLyte-647, anesthetized mice were transcardially perfused with heparinized PBS followed by 4% PFA. After 24 hours post-fixation in 4% PFA, uDISCO was performed as previously described (*14*). Subsequently, the transparent mouse heads were imaged at a z-interval of 10 µm using a LaVision BioTec Ultramicroscope II light-sheet microscope with an Olympus MV PLAPO 1x air-objective and MVX10 0.63x zoom body. Amira Software 6.3.0 was used for tracing and 3D-reconstruction of the hAβ signal in eye and optical nerve.

**Intravascular tracer administration and quantification**
A 10 µl volume of isotonic saline containing the $K^+$-analogue $^{83}Rb^+$ (0.1 µCi/µl, Perkin Elmer) was injected into the right internal carotid artery along with $^{3}H$-mannitol (0.1 µCi/µl, American Radiolabeled Chemicals) over two sec through a catheter connected to a syringe pump (Harvard Apparatus). The ipsilateral eye and optic nerve were quickly dissected, and the nerve was separated into proximal and distal parts of equal length. Radioactivity of these parts was measured using a Multipurpose Scintillation Counter (Beckman Coulter) following overnight solubilization at 45°C in Soluene (PerkinElmer) and scintillation counting with Hionic Fluor cocktail (Perkin Elmer). $^{86}Rb^+$ counts were plotted after subtraction of counts in the contralateral eye and optic nerve. FITC-cadaverine (100 µl, 0.5%, Thermo Fisher Scientific) was also administered to anesthetized mice through a catheter inserted through the external carotid artery into the right internal carotid artery, and tissue was harvested 10 min later. Fresh, unfixed eye and nerve were immediately imaged with controls from a naïve mouse under a stereofluorescent microscope (MVX10, Olympus) using a digital camera (C11440, Hamamatsu) controlled by MetaMorph software (Molecular Devices). Retina flat-mount images were collected using a confocal microscope (FV1000, Olympus) following fixation.

**Imaging and quantification of tracer transport**

For macroscopic imaging of tracer transport, the freshly harvested nerves were imaged using the stereofluorescent microscope described above. The fluorescent light intensity and exposure were kept constant across all groups. Following intravitreal injection, the ipsi- and contralateral nerves were imaged simultaneously. After subtracting the mean fluorescent intensity of the control nerve, the fluorescent intensity profile of the tracer-exposed mouse or rat optic nerve was quantified at 3.09 µm or 6.76 µm steps, respectively, using a custom-made MATLAB (MathWorks) script. CD-1 mouse optic nerves were imaged with an AxioImager M1 epifluorescence deconvolution microscope (Carl Zeiss AG) and quantified at 2.13 µm steps. Fluorescence as a function of distance from the eye was then used for further quantifications. Total fluorescence of hAβ or dextran transport was obtained by calculating the trapezoidal numerical integration of the function. Peak intensity was selected as the highest value of the fluorescent intensity profile. Movement distance was defined as the length of the wavefront of signal in the experimental nerve that exceeded by three standard deviations the mean intensity of the control nerves. A confocal scanning microscope attached to an inverted microscope (IX81, Olympus) controlled by Olympus Fluoview 500 software was used to map the cellular and subcellular hAβ tracer distribution using the whole-mount preparations or 20 µm cryosections.

**Quantitative immunohistochemistry**

Mice were anesthetized and perfused transcardially with 4% ice-cold PFA (Sigma Aldrich). Wheat germ agglutinin (lectin) tagged with AF-647 (Fisher Scientific W32466) was diluted in PBS to a final concentration of 12.5 µg/ml to label the blood vessel lumen in select experiments. Mouse optic nerve along with the orbit was post-fixed in 4% PFA for two hours. Serial 20 µm tissue sections were obtained using a cryostat (Leica CM190) following sucrose gradients that ended with 30% sucrose. Sections were incubated overnight at 4°C with a single or a combination of the following primary antibodies: rat anti-LYVE1 (ALY7) (1:250, eBioscience 14-0443-83), rabbit anti-RBPMS 1:250, GeneTex118619), mouse anti-TUJ1 (1:500 Covance MMS-435P), rabbit anti-APP (1:500, Abcam ab15272) Syrian hamster anti-podoplanin (1:100 eBioscience, clone 8.1.1), rabbit anti-AQP4 (1:500, Millipore AB3594), goat anti-mouse VEGFR3 (1:500, R&D Systems AF743), goat anti-Olig2 (1:500, R&D Systems AF2418). The primary antibodies were detected with appropriate AF-488-, 594-, 647-conjugated or Cy2-, Cy3-, Cy5-conjugated secondary antibodies (1:500, Molecular Probes/Jackson ImmunoResearch) following two hours of incubation at room temperature. Prolong Gold containing DAPI (Invitrogen, P16931) was used for mounting the sections.

**Intraocular and intracranial pressure measurement and manipulation**

Intraocular pressure (IOP) of the rodent's eye was measured by a rebound tonometer (Icare ® TONOLAB) following the manufacturer's instructions. Briefly, anesthetized rodents were placed in a prone position. Five to ten measurements at five min following induction of anesthesia were taken for each rodent to get an average baseline value. To study whether intravitreal injection induced an elevation of IOP above physiological baseline, we monitored IOP and compared it to measurements in animals with a sham surgery in which only the fluid infusion was withheld. Intracranial pressure (ICP) was monitored through a CM catheter, which was sealed to the skull with dental cement (Stoelting Co). The pressure signal was acquired at 10,000 Hz, digitized, and recorded continuously for the duration of the experiment with a DigiData 1332A and PClampex9.2 software (Axon Instruments) and then reduced to a 100 Hz sampling rate. To elevate the mouse's ICP, we infused artificial CSF (aCSF) into the CM by a Hamilton syringe connected to syringe pump (Harvard Apparatus). Artificial CSF contained the following (in mM): 155 NaCl, 3.5 KCl, 1 $CaCl_2$, 1 $MgCl_2$, and 2 $NaH_2PO_4$, pH 7.4, 300 mOsm. To lower the ICP, we actively withdrew CSF via a surgical opening in the CM. ICP was monitored in real-time during ICP manipulation. Intravitreal tracer injection was initiated five min after ICP had stabilized.

**Electron microscopy**

D2-control and DBA/2J mice were fixed by transcardial perfusion with 0.5% glutaraldehyde and 4% formaldehyde in 0.1 M phosphate buffer (pH 7.4, 9 ml/min for 25 min). The eye and optic nerve were post-fixed for two hours with 4% PFA, 1% glutaraldehyde in 0.1 M PBS before 50 µm coronal sections were dissected and embedded in Durcupan (Sigma-Aldrich). Ultrathin sections were cut, counterstained, and examined. Digital images were recorded at a nominal magnification of 4200x.

**Real-time tetramethylammonium ($TMA^+$) iontophoresis**

Optic nerves from anesthetized mice were quickly dissected following decapitation and immersed for 60 min in oxygenated aCSF at room temperature. Microelectrodes were fabricated from double-barreled theta-glass using a tetraphenylborate-based ion exchanger. The tetramethylammonium ($TMA^+$) barrel was backfilled with 150 mM $TMA^+$-chloride, while the reference barrel contained 150 mM NaCl and 10 µM AF-594. A series of currents of 20 nA, 40 nA, 80 nA, and 100 nA were applied by a dual-channel microelectrode pre-amplifier. The electrode tips were imaged after insertion into the nerve using two-photon excitation to measure the gap between the electrodes (typically ~150 µm). The $TMA^+$ signal was calculated by subtracting the voltage measured by the reference barrel from the voltage measured by the ion-detecting barrel. We used the Nikolsky equation for calibration of the $TMA^+$ electrodes based on measurements obtained in electrodes containing standards of 0.5, 1, 2, 4, and 8 mM $TMA^+$-chloride in 150 mM NaCl. The $TMA^+$ measurements were acquired relative to similar recordings obtained in 0.3% agarose prepared from a solution containing 0.5 mM $TMA^+$ and 150 mM NaCl. In-house software coded in MATLAB ('Walter', developed by C. Nicholson) was used to calculate $\alpha$ and $\lambda$ coefficients of the Nikolsky equation.

**Statistical analysis and data availability**

Statistical analysis was performed with the aid of GraphPad Prism 6.0c (GraphPad Software). Definite outlier (Q=0.1%) was removed by a ROUT method. For samples with normal distributions, we used a two-tailed *t*-test for comparisons between two groups and ANOVA for more than two groups. ANOVA was followed by Tukey's or Dunn's multiple comparisons *post hoc* test when necessary. If data did not follow a normal distribution, as indicated by a Kolmogorov Smirnov test, we used a nonparametric Mann-Whitney test or Wilcoxon matched-pairs signed rank test for comparisons between two groups or Kruskal–Wallis for comparisons among more than two groups, respectively. A Dunn's test was utilized for *post hoc* multiple comparison.

Probability values <0.05 were deemed significant. All values are expressed as the mean ± SEM.

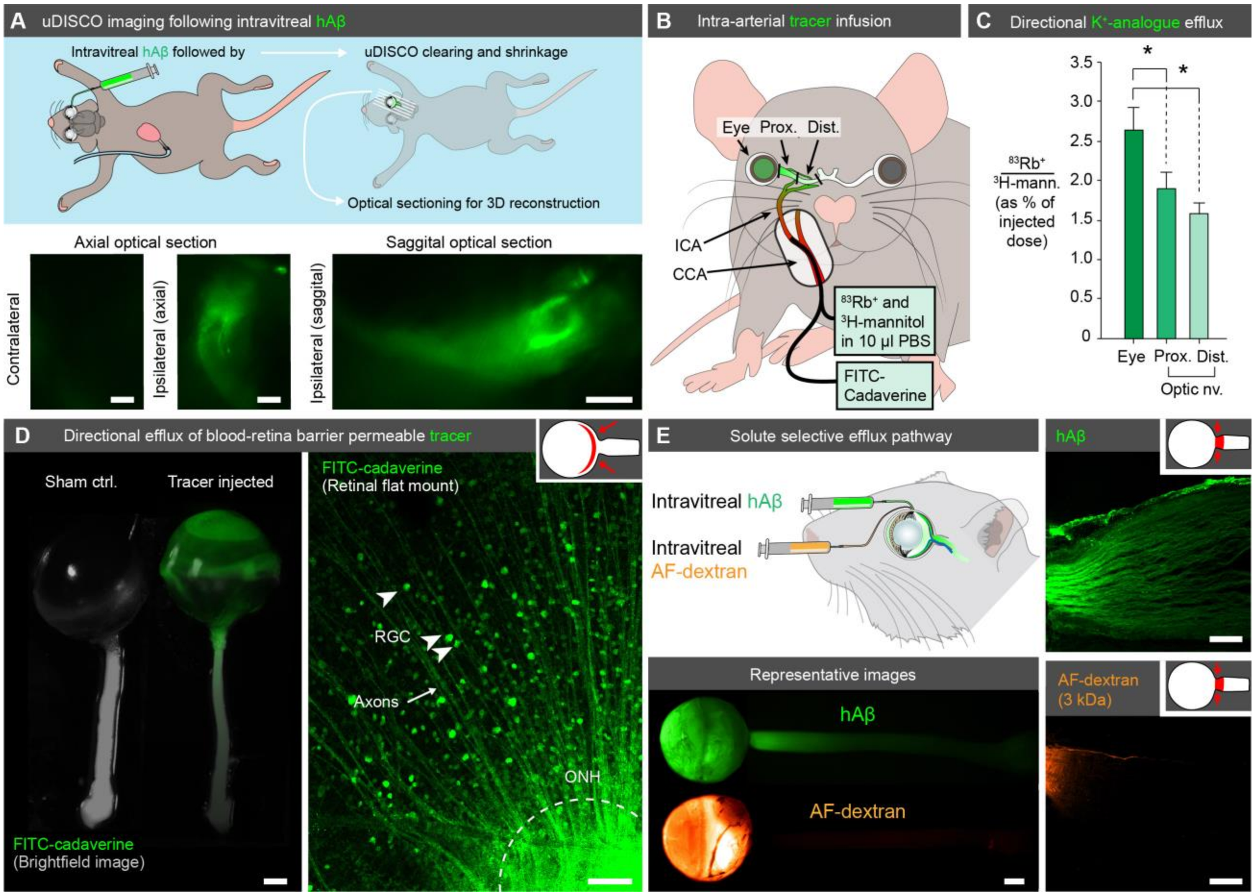


**Fig. S1. Characterization of additional tracers and tracer delivery routes.** (**A**) Top: schematic showing hAβ injection followed by uDISCO whole-mouse clearing and shrinkage, along with optical sectioning using light sheet microscopy. Bottom: representative maximum projections acquired in axial and sagittal plane ipsilateral and contralateral to the hAβ injection (scale: 500 µm). (**B**) Schematic showing non-invasive tracer delivery via injection into the internal carotid artery (ICA). (**C**) Quantification of $^{86}Rb^{+}$/$^{3}H$-mannitol radioactivity in the ipsilateral eye and optic nerve following ICA injection shows directional flow of the $K^{+}$-analogue from the eye to the optic nerve (mean±SEM, n=10, *$p<0.05$, one-way ANOVA followed by Dunnett's *post hoc* test). (**D**) Left-panel: macroscopic imaging following FITC-cadaverine ICA injection showing ipsilateral eye and optic nerve (right) compared to that from a naïve mouse (left). FITC signal (green) is superimposed on bright field (scale: 1 mm). Right-panel: confocal image of the mouse retina following ICA injection of FITC-cadaverine showing retinal ganglion cells (RGC) and their axons taking up the blood-ocular barrier-permeable tracer (scale: 100 µm). (**E**) Fluorescent macroscopic (scale: 500 µm) or confocal images (scale: 100 µm) of the eye and optic nerve following intravitreal injection of either the axonal-transportable tracer hAβ or the non-transportable tracer AF-dextran (3 kDa).

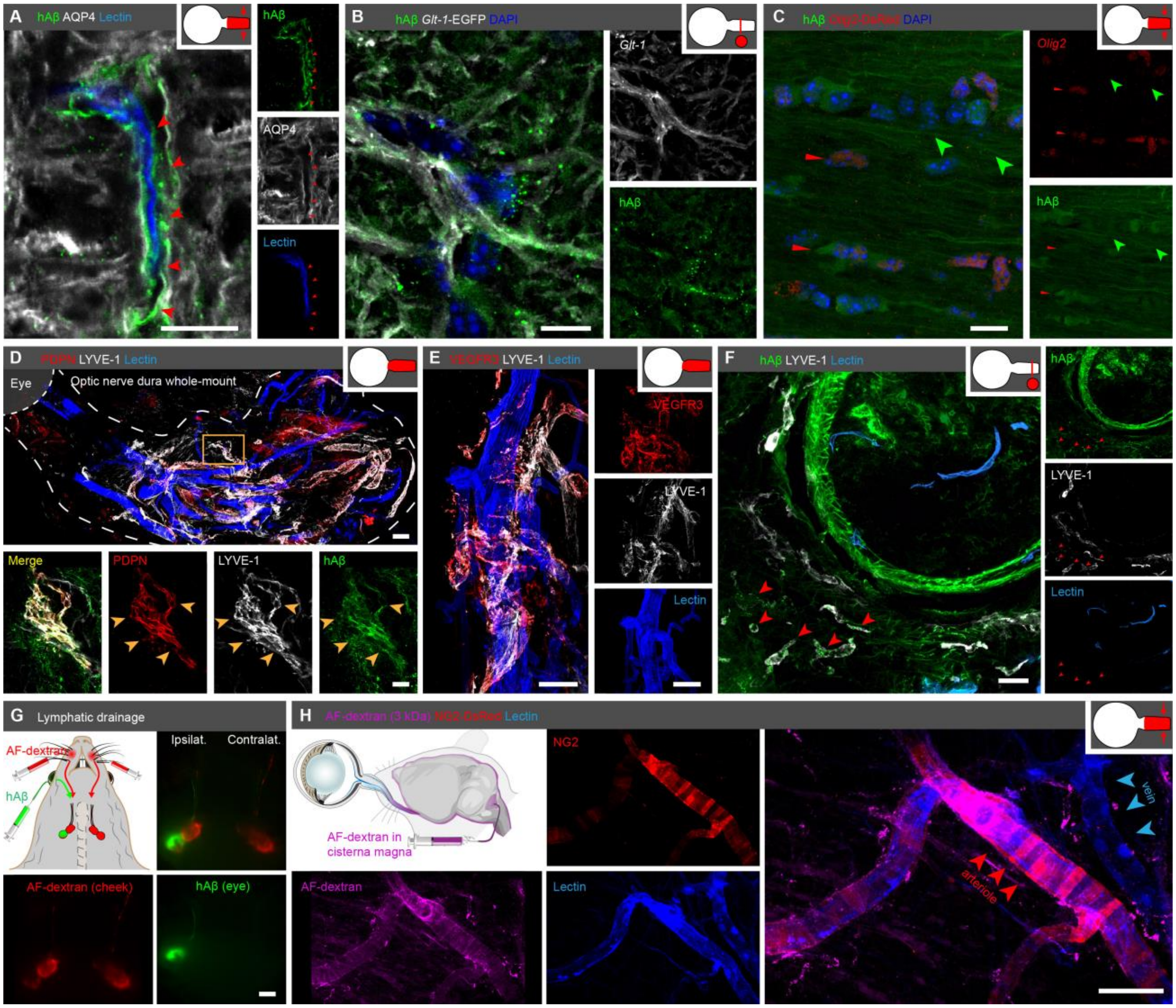


**Fig. S2. Detailed analysis of ocular glymphatic clearance route.** (**A**) Confocal image showing preferential hAβ tracer movement along the AQP4-outlined perivascular space (red arrowheads, scale: 10 µm). (**B-C**) Confocal images showing sparse uptake of hAβ tracer by *Glt-1*-eGFP- and Olig2-positive cells (scale: 10 µm; red arrowheads indicate Olig2-positive soma; green arrowheads indicate hAβ signal in axons). (**D**) Whole-mount of dura stripped from the optic nerve following intravitreal hAβ, showing LYVE-1-, PDPN-, and hAβ-positive vessel-like structures (yellow arrowheads), devoid of intravascular lectin (scale: 100 µm in top image, 20 µm in bottom images). (**E**) Dural whole-mount showing LYVE-1- and VEGFR3-positive structures, devoid of intravascular lectin (scale: 100 µm). (**F**) Immuno-detection of LYVE-1 in cross-section of optic nerve following hAβ tracer intravitreal injection, showing co-labeling of LYVE-1-positive, intravascular lectin-negative structures (red arrowheads) with hAβ tracer in the dura and orbital tissue surrounding the optic nerve (scale: 20 µm). (**G**) Schematic and macroscopic imaging of cervical lymph nodes following intravitreal hAβ injection (scale: 1 mm). (**H**) Confocal image of the optic nerve from an NG2-DsRed mouse following cisterna magna (CM) injection shows AF-dextran tracer accumulated preferentially in the periarterial space rather than the perivenous space (scale: 50 µm).

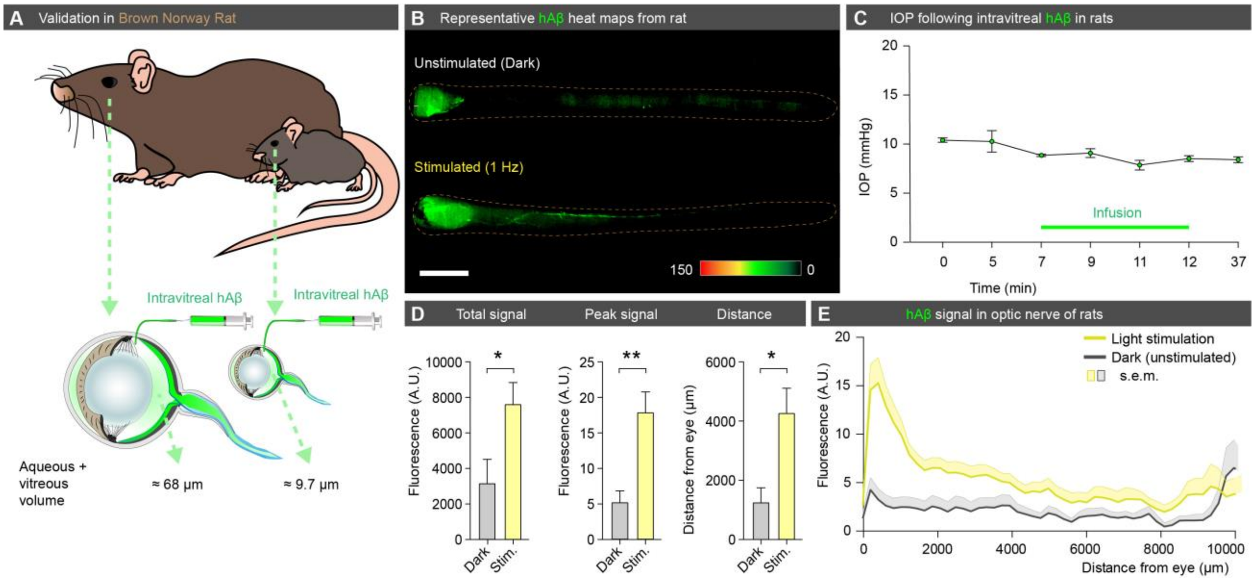


**Fig. S3. Validation of tracer delivery model in rats.** (**A**) Schematic showing intravitreal hAβ infusion in Brown Norway rat, which has approximately seven times the intraocular fluid volume of a C57BL/6J mouse. (**B**) Representative background-subtracted heat maps from rats kept in darkness or subjected to 1 Hz light stimulation following intravitreal hAβ infusion demonstrating a similar pattern to that of a mouse (scale: 1 mm). (**C**) IOP measured by rebound tonometry during the hAβ infusion in rats (mean±SEM, n=3). (**D**) Total hAβ signal, peak hAβ signal, and distance travelled for unstimulated and stimulated (1 Hz) rat eyes (mean±SEM, n=5-7, *$p<0.05$, **$p<0.01$, unpaired two-tailed $t$-test). (**E**) Averaged fluorescent intensity profile of hAβ tracer distribution along rat optic nerve (mean±SEM, n=5-7).

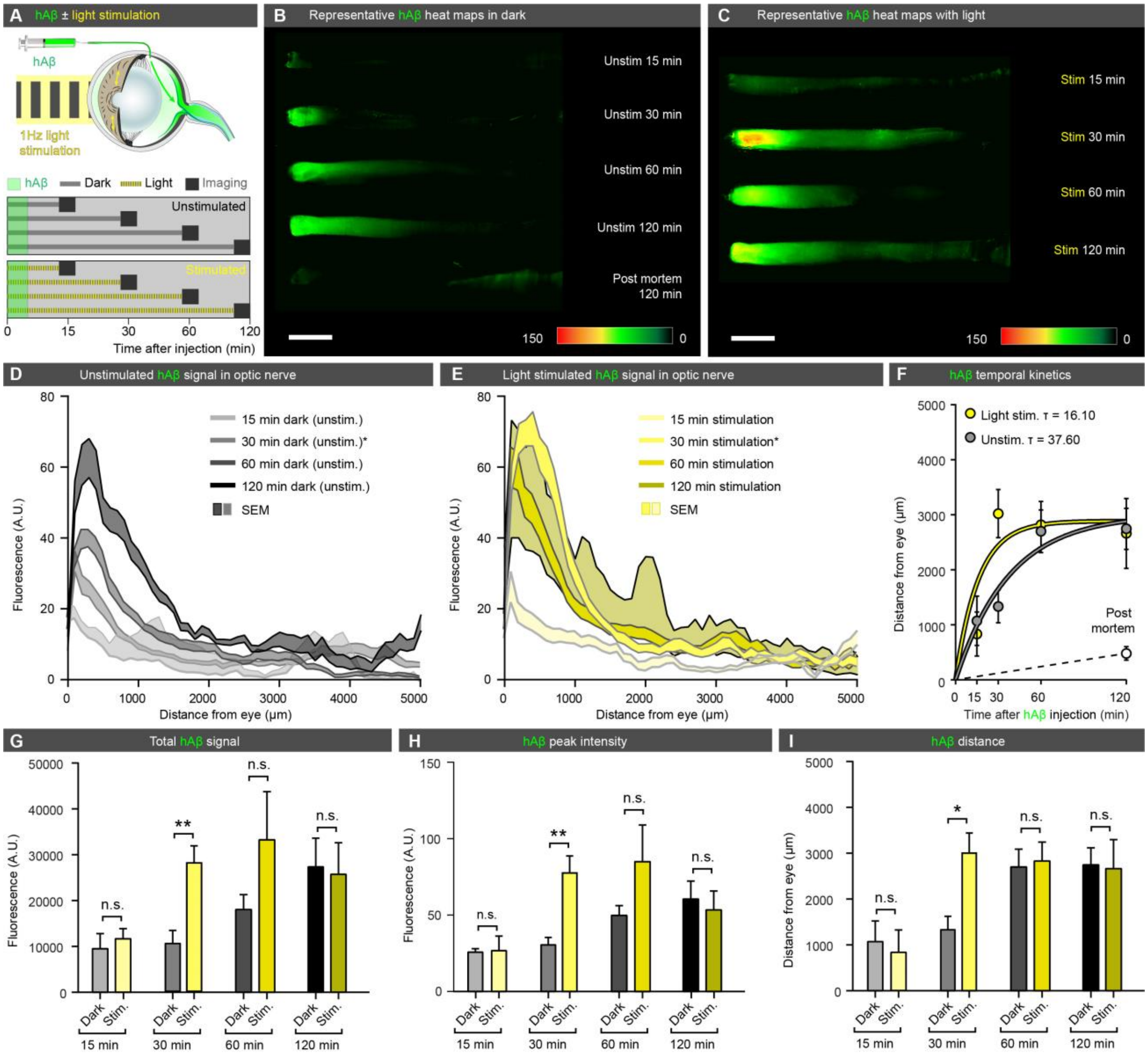


**Fig. S4. Light stimulation accelerates hAβ tracer movement along the optic nerve.** (**A**) Mice were either subjected to dark or 1 Hz light stimulation (100 ms duration, 5 lumens) and imaged 15, 30, 60, or 120 min after intravitreal hAβ injection. (**B-C**) Representative background-subtracted heat maps of optic nerves following hAβ tracer injection demonstrate that light stimulation accelerates hAβ tracer clearance via the optic nerve. Almost no hAβ tracer movement is seen postmortem (scale: 1 mm). (**D-E**) Averaged fluorescent intensity profiles of hAβ tracer distribution along the optic nerve in dark unstimulated and light-stimulated groups at 15-, 30-, 60-, and 120-minute time points (mean±SEM, n=4-8). (**F**) Pseudo-kinetics of the hAβ tracer movement shown by fitting the distance and experiment time to a one-phase decay curve (mean±SEM, n=4-8). Quantification of the time constant (tau, **τ**) shows light stimulation induces a two-fold acceleration in tracer movement. (**G-I**) Total hAβ tracer signal, peak intensity, and distance travelled in the optic nerve for all the groups. Light stimulation causes significantly increased hAβ tracer transport in the optic nerve 30 min after intravitreal injection (mean±SEM**,** n=4-8, *$p$<0.05, **$p$<0.01, n.s. $p$=0.193-0.946, unpaired two-tailed $t$-test).

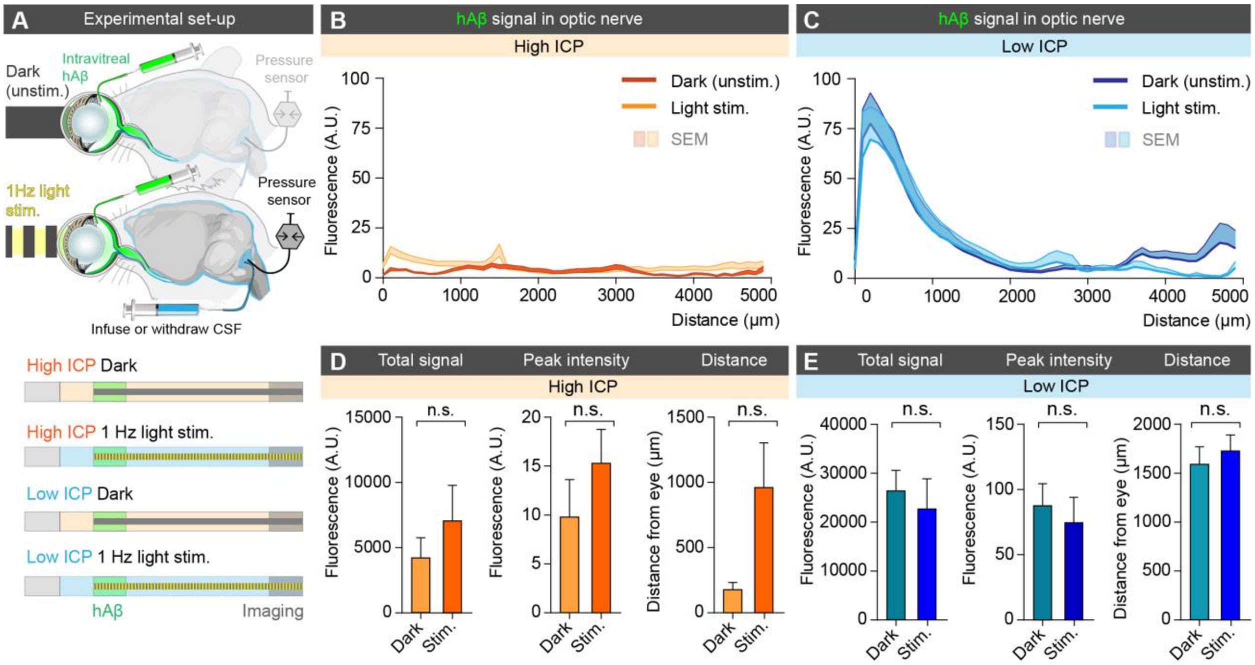


**Fig. S5. Influence of the translaminar pressure gradient on stimulated and unstimulated tracer movement.** (**A**) Schematic of experimental design used for intravitreal injection with concurrent manipulation of intracranial pressure (ICP) and exposure to 1 Hz light stimulation or dark (unstimulated). (**B-C**) Averaged fluorescent intensity profiles of hAβ tracer distribution along the optic nerve in unstimulated and 1 Hz light-stimulated groups under high and low ICP conditions (mean±SEM, n=6). (**D-E**) Total hAβ tracer signal, peak intensity, and distance of hAβ tracer transport in the optic nerve following intravitreal injection in either high ICP condition or low ICP condition (mean±SEM, n=6, n.s. $p$=0.104-0.627, unpaired two-tailed $t$-test).

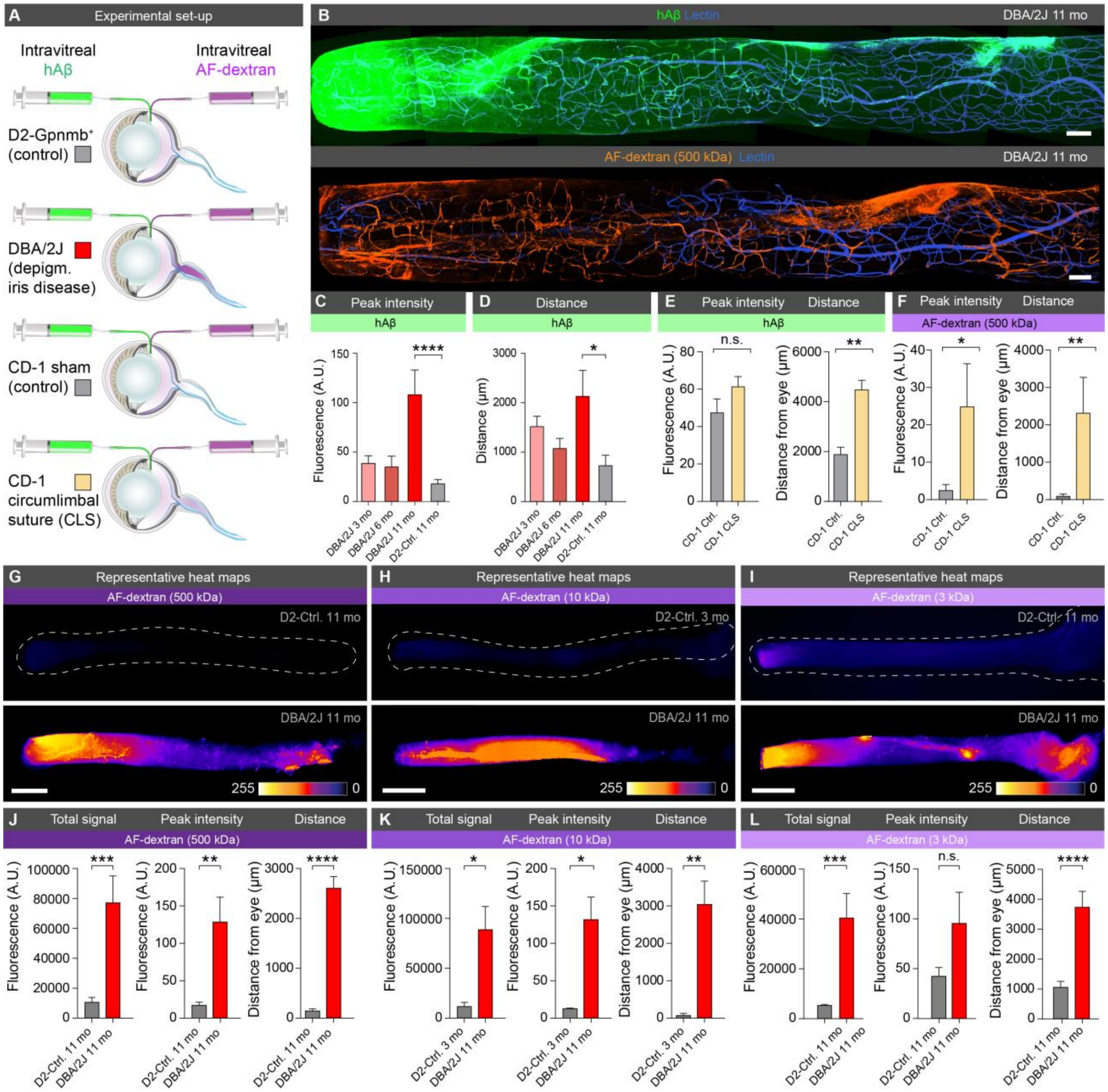


**Fig. S6. Murine glaucoma models reveal defects in the lamina barrier allowing escape of large intraocular tracers.** (**A**) Schematic of the experimental set-up for intravitreal injection of different size AF-dextran tracers (3, 10, 500 kDa) and hAβ tracer in DBA/2J and chronic circumlimbal suture (CLS) glaucoma models. (**B**) Representative confocal images of the ipsilateral optic nerve following intravitreal injection of hAβ tracer (top-panel) and AF-dextran (bottom-panel) in old DBA/2J mice (scale: 100 µm). (**C-F**) Peak intensity and distance of hAβ and AF-dextran (500 kDa) tracer transport in the optic nerve in 11-month-old DBA/2J, age-matched D2-control, as well as young and mid-age DBA/2J and CLS mice along with CD-1 controls (mean±SEM, n=6-12, *$p$<0.05, **$p$<0.01, ****$p$<0.0001, n.s. $p$=0.1255, Kruskal-Wallis test followed by Dunn's *post hoc* test or unpaired two-tailed $t$-test, or Mann-Whitney test). (**G-I**) Representative background-subtracted heat maps of optic nerve following AF-dextran (3, 10, 500 kDa) tracer injection in D2-control and old DBA/2J mice (scale: 500 µm). (**J-L**) Total hAβ tracer signal, peak intensity, and distance of AF-dextran tracer transport along the optic nerve for 500, 10, and 3 kDa AF-dextran (mean±SEM, n=6-9 [500 kDa groups], n=4-8 [10 kDa groups], n=5-6 [3 kDa groups], $p$=0.0917 for peak signal 3 kDa AF-dextran DBA/2J vs. control, *$p<0.05$, **$p<0.01$, ***$p<0.001$, n.s. $p$=0.069, unpaired two-tailed $t$-test).

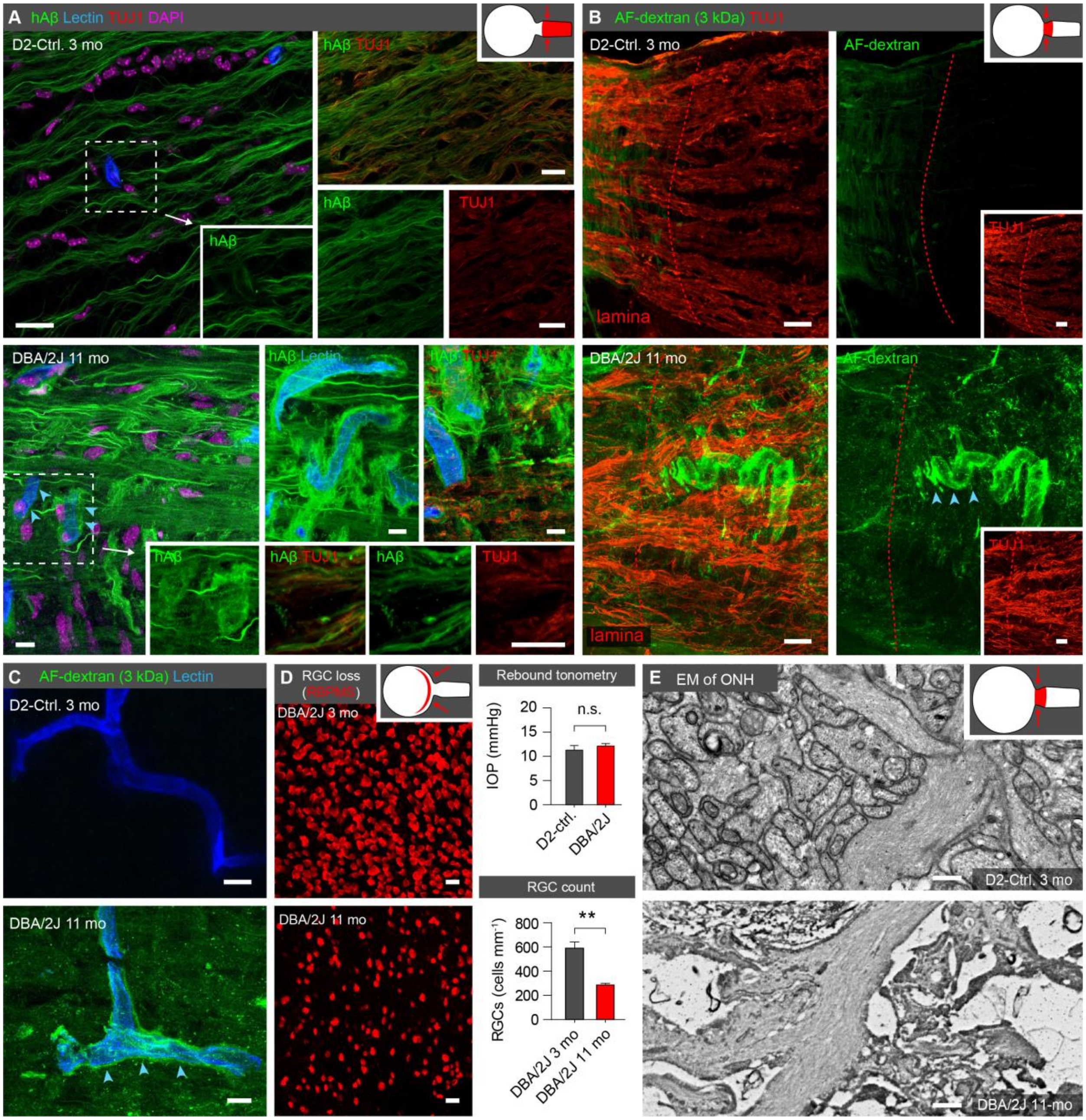


**Fig. S7. Tracer efflux pattern in murine glaucoma model indicates leaky lamina that may allow bypass of intra-axonal transport via extracellular route.** (**A**) Representative confocal images of the ipsilateral optic nerve following intravitreal hAβ tracer injection from D2-control (top) and 11-month-old DBA/2J mouse (bottom). Note the mixture of axonal, extracellular, and perivascular hAβ tracer transport (blue arrowheads: perivascular transport, scale: 20 µm). (**B**) Confocal images of the optic nerve head stained for axonal marker TUJ1 after intravitreal injection of AF-dextran (3 kDa) in D2-control (top) and old DBA/2J mice (bottom). Note that the dextran tracer does not penetrate the lamina barrier (red dotted line) in control mice, while old DBA/2J mice show extensive dextran leakage into the retrolaminar optic nerve (blue arrowheads: perivascular tracer, scale: 20 µm). (**C**) Higher magnification confocal image showing dextran cannot penetrate the glial barrier in control mice, while dextran tracer in old DBA/2J mice shows a predominantly extra-axonal and perivenous accumulation (blue arrowheads: perivascular tracer, scale: 10 µm). (**D**) Left-panel: representative confocal images of retina flat-mount demonstrating a loss of RGCs in old DBA/2J mice compared to young DBA/2J animals (scale: 20 µm). Right-panel: quantification of IOP and in age-matched young D2-control and DBA/2J mice as well as RGC number in young DBA/2J vs.old DBA/2J mice (mean±SEM, n=11-33 for IOP and n=9-10 for RGC count, $^{**}p<0.01$, n.s. $p=0.436$ unpaired two-tailed *t*-test). (**E**) Representative electron micrographs from the glial lamina region of young D2-control (top-panel) and old DBA/2J mice (bottom-panel) (scale: 0.5 µm) shown here without pseudo-coloring as compared to Fig. 4i.

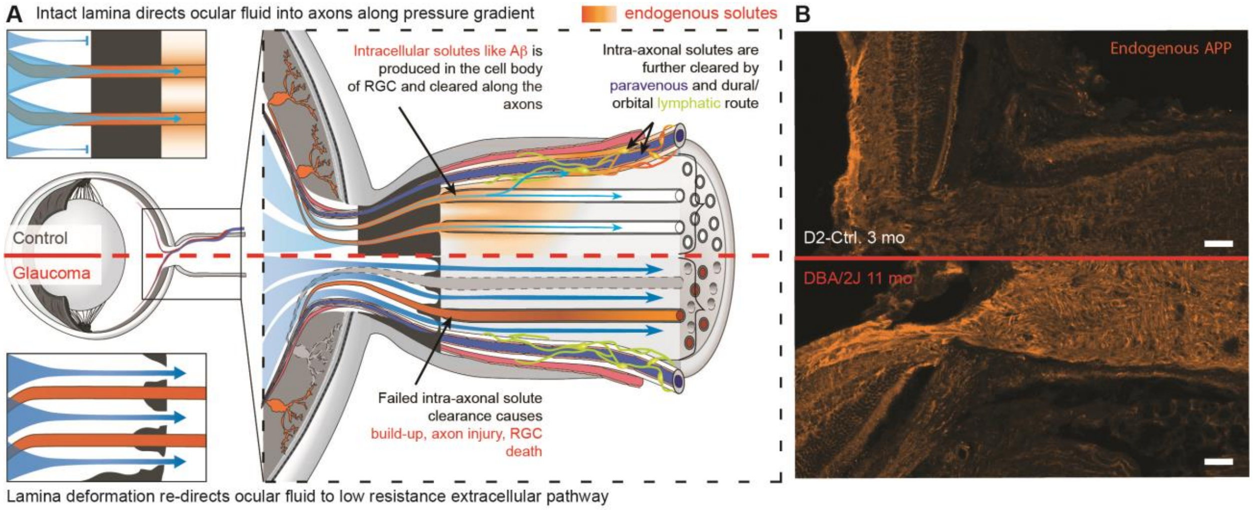


**Fig. S8. Schematic model of the ocular glymphatic clearance system and its dysfunction in a murine model of glaucoma.** (**A**) In control mice (top), IOP drives ocular fluid (IOF, blue) into RGCs axons prior to the glial cell barrier of lamina cribrosa (black). Intra-axonal transport of solutes (orange) across the lamina is accelerated by IOF uptake by RGC axons. After passing the glial lamina, hAβ is released into the extracellular space, disperses into the perivenous space, and is eventually cleared by dural/orbital lymphatic vessels. Insert illustrates how IOP in combination with the glial lamina barrier drives IOF into the unmyelinated RGC axons. In old DBA/2J mice (bottom), the barrier function of the glial lamina is lost, resulting in IOF passing directly into the extracellular space of the optic nerve via defects in the lamina, thus bypassing the intra-axonal compartment. The reduction of intra-axonal flow results in accumulation of potentially harmful solutes, such as Aβ (dark orange axons), and axonal degeneration (grey axons). Insert shows IOF bypassing the glial lamina through openings in the barrier. (**B**) Depicts immunolabeling of endogenous (mouse) amyloid precursor protein (APP) in young (top) and old (bottom) DBA mice (scale: 20 µm).
